# Synapse Detection Efficiency in EM *Drosophila* Connectomics

**DOI:** 10.1101/2025.10.16.682869

**Authors:** Louis K. Scheffer

## Abstract

Researchers have long noted the differences in synapse count between different EM reconstructions of similar circuitry. In this paper we attempt to determine the portion of these differences that may be due to different sample preparation and imaging techniques, in particular serial-section transmission imaging (SS-TEM) compared to focused ion beam with scanning electron microscopy (FIB-SEM). To do this, we compare synapse detection in the major *Drosophila* EM reconstructions - FANC, MANC, FAFB (with original and new synapses), male CNS, BANC, and HemiBrain, plus several smaller reconstructions. We look at raw synapse counts to avoid any dependence on proofreading, and compensate insofar as possible for the confounds of sample sizes differences and different software detection efficiency. The result are estimates, per compartment and for the sample as a whole, of the number of synapses that would be visible to a skilled human observer. These are then compared across all samples, using regions which are reconstructed in common for each sample pair. We find that in almost all known cases where a volume has been reconstructed by both techniques, isotropic FIB-SEM reconstructions show more human-visible synapses than microtome sliced reconstructions, typically by more than 40%. This strongly suggests, but does not conclusively prove, that synapses are easier to see in isotropic FIB-SEM data.

## Introduction

With advances in EM reconstruction, there are now several different reconstructions of portions of the adult *Drosophila* brain. The reconstructions considered in the paper are shown in Fig 1. Here we compare synapse counts of different reconstructions.

**Figure 1:**
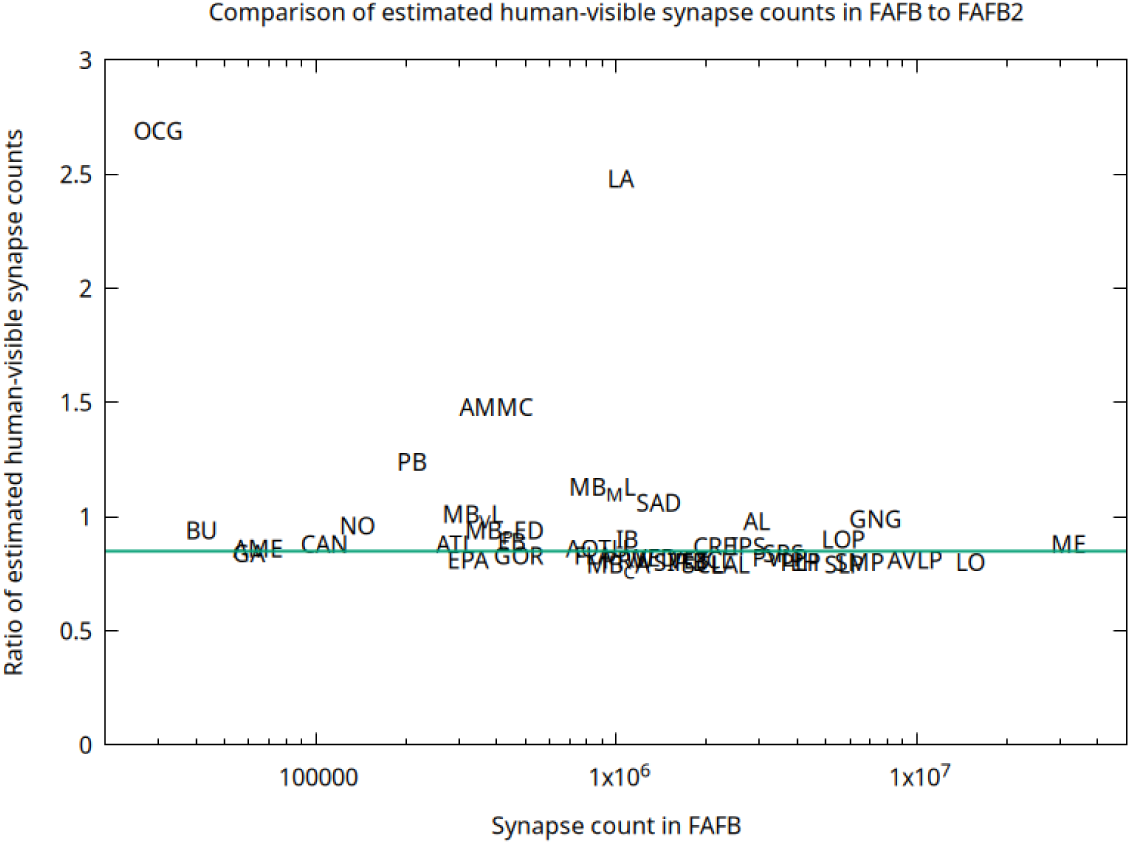
Comparison of estimated human-visible synapse counts in FAFB and FAFB2 (new synapse set), for top-level regions. Similar regions found on both sides of the brain have been combined. Data point is the center of the corresponding label. This is a scaled version of the data presented as Figure 5 of (Yu et al. 2025).

**Table 1:** **Reconstructions considered here. FANC fly^†^** was genotype y,w/w[1118]; +; P{VT025718-Gal4}attP2/P{pBI-UASC–3 x MYC–sbAPEX2–dlg-S97}18. VNC stands for Ventral Nerve Cord.

| name | year | sex | type | age | description | technology |
| --- | --- | --- | --- | --- | --- | --- |
| 1C | 2013 | F | Oregon R | 5d | 1 optic lobe column | SS-TEM |
| 7C | 2017 | F | Canton S | 5-6d | 7 optic lobe columns | FIB-SEM |
| FIB19 | 2019 | F | Conton S | 6d | Portion of optic lobe | FIB-SEM |
| Hemibrain | 2020 | F | Canton S | 5d | Half central brain | FIB-SEM |
| FANC | 2021 | F | See note<sup>†</sup> | 1-2d | Female VNC | SS-TEM |
| FAFB | 2024 | F | Canton S | 7d | Central brain | SS-TEM |
| FAFB2 | 2025 | F |  | 7d | New synapses |  |
| MANC | 2024 | M | Canton S | 5d | Male VNC | FIB-SEM |
| CNS | 2025 | M | Canton S | 5d | Brain and VNC | FIB-SEM |
| BANC | 2025 | F | Canton S | 5-6d | Brain and VNC | SS-TEM |

**Table 2:** Major *Drosophila* EM reconstructions.

| Data set | Reference |
| --- | --- |
| Hemibrain | Scheffer et al. (2020) |
| FAFB | Buhmann et al. (2021) |
| FAFB2 | Yu et al. (2025) |
| FANC | Phelps et al. (2021) |
| MANC | Takemura et al. (2023) |
| CNS | Berg et al. (2025) |
| BANC | Bates et al. (2025) |

There are two methods commonly used to take electron microscope (EM) images for neural reconstruction. The first is serial sectioning combined with transmission electron microscopy (here abbreviated SS-TEM). The sample is physically divided into thin slices, then each slice is imaged in transmission. Resolution is typically roughly 4×4×40 nm. The 40nm resolution in one dimension is determined by the minimum sample thickness that can be cut reliably.

The second method is focused ion beam milling combined with scanning electron microscopy (here abbreviated FIB-SEM). The sample is imaged, then a thin layer is abraded away by an ion beam, then another image is taken. This process is repeated until the sample is completely imaged. The resolution is typically 8×8×8 nm, limited primarily by imaging speed.

There are several reconstructions done with each of these technique. The first circuit reconstructed in the *Drosophila* adult was a single column in the optic lobe of the medulla, imaged using SS-TEM(Takemura et al. 2013). Synapses were identified by hand. The second example, and the first using FIB-SEM, was 7 columns of the hexagonal optical medulla lattice(Takemura et al. 2015). The sample was prepared similarly, specifically to facilitate comparisons, and the reconstruction was done by the same team, but machine learning was first used to identify synapse candidates. The seven column reconstructions was followed by a larger portion of the optic lobe, sample FIB19 (Shinomiya et al. 2019). Again, the sample preparation methods for FIB19 were the same as the single and 7 column samples, to facilitate comparisons among these data sets.

Another reconstruction using FIB-SEM was half of the central brain (the hemibrain)(Scheffer et al. 2020).

A female nerve cord (FANC) was imaged with SS-TEM and partially reconstructed(Phelps et al. 2021).

A male adult nerve cord (MANC) was reconstructed with FIB-SEM(Takemura et al. 2023).

A larger SS-TEM reconstruction is the Full Adult Fly Brain (here FAFB) (Dorkenwald et al. (2024) and Schlegel et al. (2024)), containing the central brain and both optic lobes.

In 2025, a new set of synapses for the FAFB data set was released. This set uses the segmentation as an aid to finding synapses, and features much better synapse recall, but somewhat less precision(Yu et al. 2025). In this paper we refer to the FAFB proofreading with the new synapses as FAFB2.

Also in 2025, a combined brain and nerve cord (BANC) of a female fly was released, using SS-TEM and 4×4×45 nm voxels(Bates et al. 2025).

Finally we have a full male central nervous system (CNS), reconstructed using FIB-SEM. The first published result of this reconstruction was the right optic lobe(Nern et al. 2024). The full reconstruction (Berg et al. 2025) contains both optic lobes, the central brain, and the ventral nerve cord.

## 1 Comparing synapse counts

One problem with comparing synapse counts is that different reconstructions use different programs to detect synapses, as there are far too many synapses to annotate by hand. This synapse detection software can vary considerably in effectiveness, depending on the technology involved and the way it is tuned. This is illustrated by the new synapse set for FAFB, which finds significantly more synapses than the original reconstruction.

One way around this is to estimate “visible” synapse counts, which are the number of synapses that could have been detected by an experienced and proficient human. This can be estimated, as each reconstruction has measured the *recall* of synapse detection, which is the fraction of all humanly visible synapses that are found by the software, and the *precision*, which is the fraction of found synapses that are real. This is typically measured by taking subvolumes of tissue scattered over the whole reconstruction, then directing human annotators to find all possible synapses in these samples. These are then compared to software results of synapse detection on the same subvolumes, to generate statistics for recall, the percentage of human-visible synapses found by the algorithm, and precision, the percent of machine-found syapses that are real. For the purpose of our analysis we need the precision and recall numbers, as shown in Table 3.

**Table 3:** Published statistics and estimated human-visible synapses. Autapses have been removed from all data sets.

| Data set | detected | recall | precision | visible |
| --- | --- | --- | --- | --- |
| Hemibrain | 64.1M | 0.80 | 0.80 | 64.1M |
| FAFB | 130.1M | 0.65 | 0.95 | 190.1M |
| FAFB2 | 187.9M | 0.95 | 0.82 | 162.2M |
| FANC | 45.1M | 0.73 | 0.71 | 43.9M |
| MANC | 74.5M | 0.80 | 0.80 | 74.5M |
| CNS | 311.8M | 0.80 | 0.80 | 311.8M |
| BANC | 213.8M | 0.95 | 0.68 | 153.0M |

These measured numbers allow us to reverse the synapse detection process, estimating how many synapses *could* have been found by humans if they had the time to do so. Remember that *precision* is the fraction of detected synapses that are real, and *recall* is the fraction of real synapses that are detected. Then

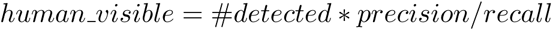

This process can be demonstrated, for example, with the two different synapse detection methods used on FAFB. Here the initial algorithm found 130M synapses, and with a precision of 0.95, 123.5M of these are real. Then with a recall of 0.65, we expect about 190M synapses that could be detected by humans. So precision is 123.5/130 = 95%, as desired, and recall is 123.5/190 = 65%, also as desired.

The second synapse detection used different methods and chose different tradeoffs, with a higher recall of 0.95 accompanied by somewhat lower precision of 0.82. This method found 188M synapses, which combined with their measured precision of 0.82 implies 154M are real. Then their recall of 0.95 implies 162M synapses could be visible to humans. Again these numbers are consistent with the reported detections, and precision and recall figures. Further, this relationship holds at the compartment level, though with more scatter. Fig. 1 shows the ratio of the estimated human-visible synapses in each top level compartment, based on the measured counts and the global recall numbers. Each compartment would fall at *Y* = 0.85 if the the global precision and recall number applied uniformly to all compartments. Even though the new synapse set detects 46% more synapses, the estimated human-visible synapse counts are much closer, except for a few smaller compartments (such as OCG and LA) where the new synapse prediction finds many more synapses (likely because of new and improved ground truth for those compartments).

The visible-synapse compensation using recall also appears to work (though with significant scatter) on the individual neuron level, as shown in Fig. 2. This figure compares two different synapse detection algorithms run on FAFB sample. The estimated number of human-visible synapses is computed for each. If the compensation for synapse recall was perfect, the two estimated curves would be identical, since the sample is the same. Although not identical, the means are much closer for the human-visible estimates (8% different), compared to the raw numbers of detected synapses (34% different).

**Figure 2:**
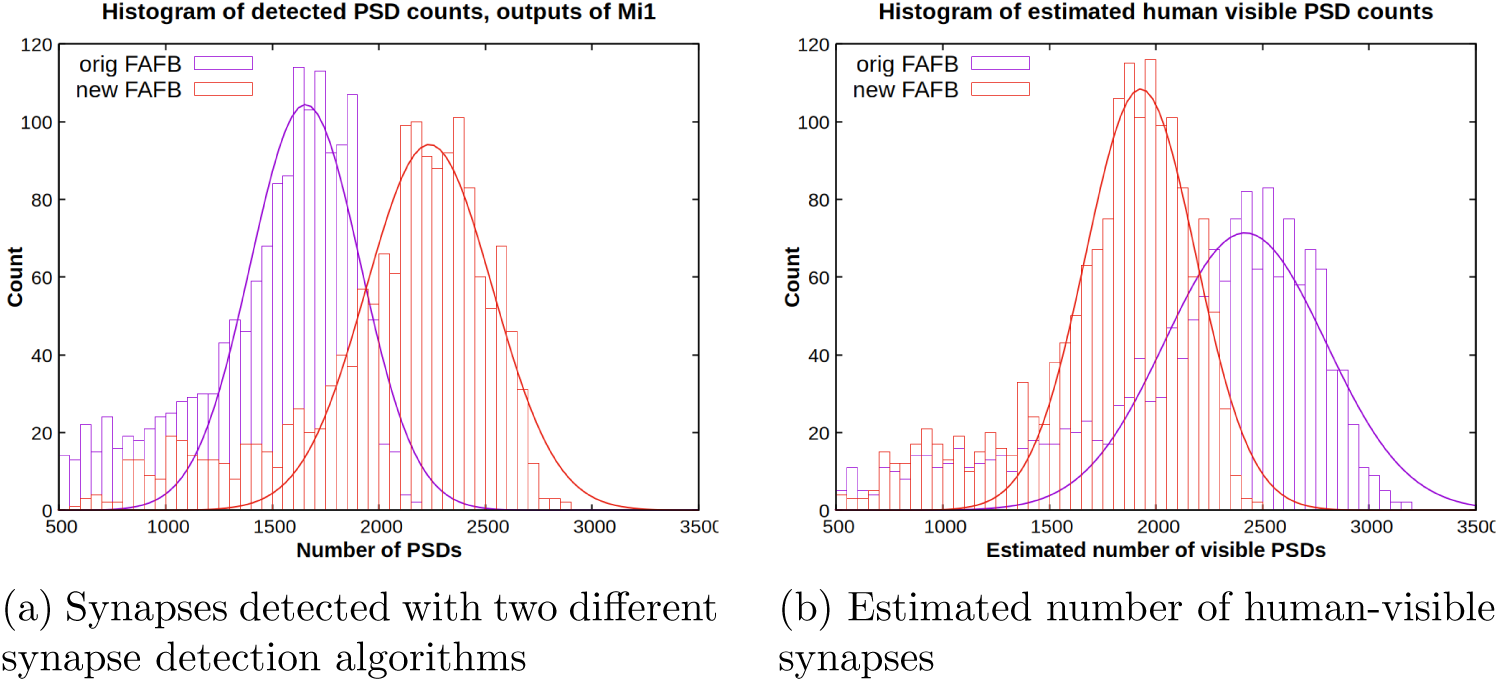
Comparison of two different synapse detection algorithms (FAFB and FAFB2) run on the same image data. Displayed are histograms of PSD counts of outputs of all Mi1 cells in the sample. The estimated number of human-visible synapses are more consistent than the detected counts.

### 1.1 Comparing estimated synapse counts

Given synapse number per compartment, comparing synapse counts in the central brain of the largest reconstructions - Hemibrain, FAFB, male CNS, and BANC - is straightforward. These samples have all been divided by their respective authors into regions of interest (ROIs), with consistent names across reconstructions. For our comparison, we sum over all compartments fully captured in both samples. This requirement of full compartments is mostly relevent for the hemibrain, where we use the abbreviation HBI, for hemibrain (internal), meaning only compartments fully internal to the sample.

For samples including the VNC (Ventral Nerve Cord), however, compartment names are only consistent between CNS and MANC, and are different from those in BANC and FANC. Correspondence tables (Stuart Berg, personal communication) must be applied, and are shown in Appendix 3. These samples are then compared using the same technique as full brains.

The pairwise results of these comparisons is shown in Table 4.

**Table 4:** Ratios of estimates of human-visible synapse counts in overlapping regions of reconstructions. . Order within each technology is arbitrary but is arranged so that ratios are in general greater than one for ease of reading. Ratios in lower left are inverses of those in upper right, and are not shown.

|  | Imaging | SS-TEM |  |  |  | FIB-SEM |  |  |
| --- | --- | --- | --- | --- | --- | --- | --- | --- |
|  | Name | BANC | FANC | FAFB | FAFB2 | HBI | CNS | MANC |
| TEM | BANC | 1 | 1.13 | 1.64 | 1.40 | 1.66 | 1.95 | 1.94 |
|  | FANC |  | 1 | — | — | — | 1.45 | 1.73 |
|  | FAFB |  |  | 1 | 0.87 | 0.89 | 1.26 | — |
|  | FAFB2 |  |  |  | 1 | 1.10 | 1.45 | — |
| FIB | HBI |  |  |  |  | 1 | 1.34 | — |
|  | CNS |  |  |  |  |  | 1 | 1.18 |
|  | MANC |  |  |  |  |  |  | 1 |

The full brain synapse ratios are reproduced, though with more scatter, when the comparison is done at the level of brain regions. An example is the comparison for all compartments in common between FAFB2 and the male CNS. The global average is 1.49, and the individual compartments are centered around this value, though with considerable variation. See Figure 3.

**Figure 3:**
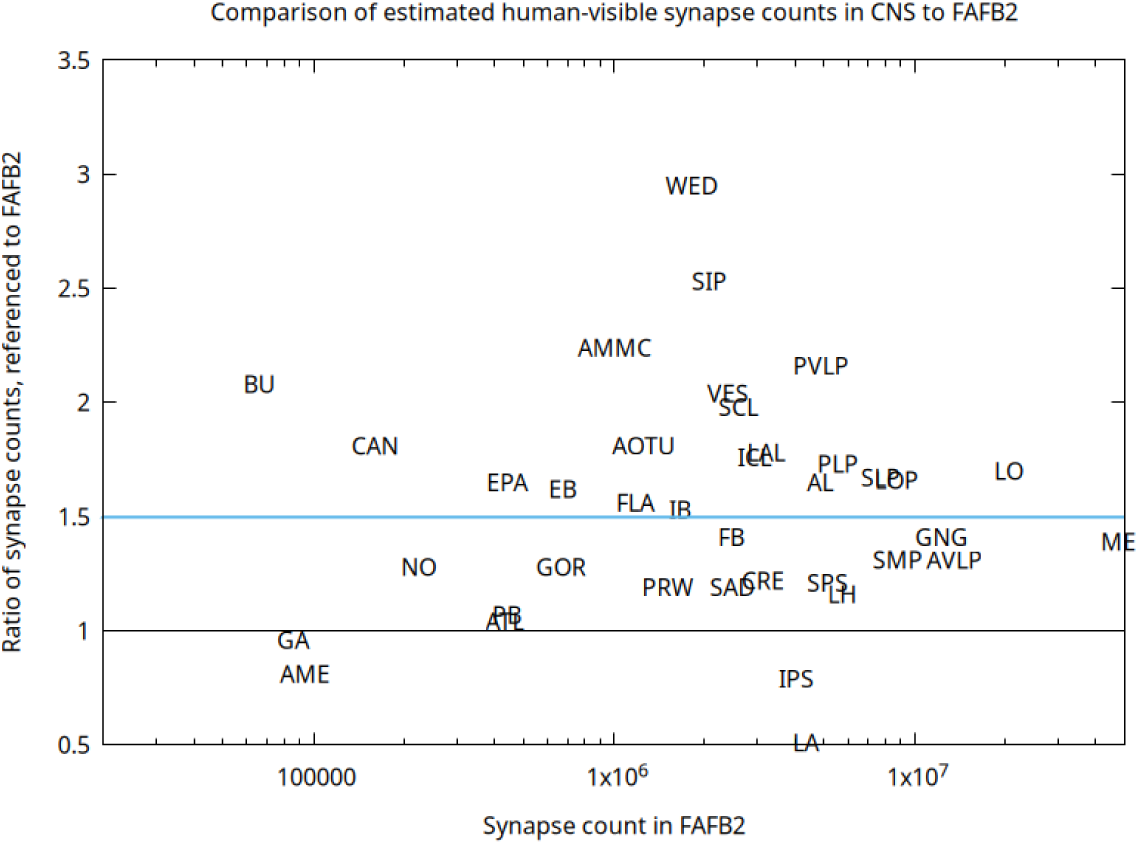
Comparison of estimates of human-visible synapse counts in CNS and FAFB2 (new synapse set), for top-level regions found in both. Similar regions found on both sides of the brain have been combined. Data point is the center of the corresponding label.

Comparisons involving the older samples - 1C, 7C, and FIB19 - are more complicated. These samples are much smaller, and have many neurons that extend out of the volume and are otherwise incompletely reconstructed. Compartment and full sample size comparisons make little sense for these sample, so only neuron level comparisons are available. Even in this case, considerable care is needed to select neurons likely to be complete. For comparison between the single column and seven column reconstructions, where these problems are particularly acute, we rely on the analysis in Takemura et al. (2015), supplementary material. For comparisons using FIB-19, we use only the central 7 columns of the medulla, those that are most completely reconstructed.

## 2 Sizes of samples

Flies vary in size, which could account for at least some of the differences in synapse count. To compare sample sizes, we divide each sample into 2 micron cubes, then find (for each neuropil/compartment) the number of cubes touched by the synapses marked with that neuropil name. For an overall size comparison for each sample pair, we sum the volume over all common compartments. This comparison is of course subject to variation in how compartments are defined, but seems to be relatively consistent over full samples (with the exception of FAFB2, which gives much larger compartment volumes that FAFB, which is odd since it’s the same sample. In this case we use the FAFB volumes, which seem more consistent with other samples.). The result is the size comparison table below.

**Table 5:** Approximate sizes ratios of reconstructions. . Order is same as the previous table. Ratios in lower left are inverses of those in upper right, and are not shown.

| Name | FANC | BANC | FAFB | FAFB2 | HBI | CNS | MANC |
| --- | --- | --- | --- | --- | --- | --- | --- |
| BANC | 1 | 1.13 | 1.15 | 1.15 | 1.32 | 1.09 | 1.10 |
| FANC |  | 1 | – | – | – | 0.97 | 0.98 |
| FAFB |  |  | 1 | 1 | 1.08 | 0.93 | – |
| FAFB2 |  |  |  | 1 | 1.08 | 0.93 | – |
| HBI |  |  |  |  | 1 | 0.82 | – |
| CNS |  |  |  |  |  | 1 | 1.02 |
| MANC |  |  |  |  |  |  | 1 |

But are the coordinates in each synapse file accurate? There are a number of transformations between the living fly and the images collected for reconstruction. First it is known that sample preparation changes sizes considerably, as the water is squeezed out and replaced by plastic. We naively assume this is roughly the same for all samples.

Next, each of the samples has been sliced in some fashion, which can change the size and shape of the sliced tissue samples. FAFB was sliced into 40nm layers, FANC and BANC into 45 nm layers, and the hemibrain, MANC, and the male CNS were sliced into roughly 20*µ* slabs before the FIB process. Fortunately, for three of the largest samples, FAFB, HB, and CNS, there exist pre-slicing CT scans of the samples. The CT scans are expected to be well calibrated, as the X-ray path involves no lenses - just a point source, the sample, and the detector. So the scale factor is determined by the source-to-sample and sample-to-detector distances, which are well measured. EM images, in contrast, depend on electron optics, which must themselves be calibrated.

Therefore we aligned these data sets to their CT scan images. This is not a procedure of great accuracy, as Xrays and electrons see different features, and precisely locating corresponding points in CT images is hard due to the low resolution. For FAFB we have only the image stack, not the meta-data (which has the absolute scale). We assumed the X direction (orthogonal to the knife cut) was the advertised 4 nm. Then the Z cuts come out at 41.75 nm/cut, which is reasonable. The Y direction, in the direction of the cut, looks compressed by about 15%, which is quite visible by eye and historically reasonable(Studer & Gnaegi 2000). This gives 4.62 nm/pixel in Y.

In the hemibrain it was particularly difficult to match features in the EM and CT, as there are no tissue edges (the easiest features to compare) in the EM. We get X = 8.025 nm/pxel, Y=7.56 nm/pixel, and Z = 8.45 nm/pixel. For the male CNS we get x=7.66 nm/pixel, Y=7.72 nm/pixel, and Z=8.00 nm/pixel.

When finding volumes for these three samples, we used these empirical sizes. For the remaining samples, FANC, BANC, and MANC, we use the specified pixel size without correction. The three CT scans we used have been uploaded to Zenodo, so these calibrations can be recreated from publicly available data.

As another rough test of fly size, we consider the total number of Mi1 cells in the samples with proofread optic lobes, FAFB, BANC, and CNS. This cell not believed to be sexually dimorphic, and occurs in large numbers in each sample. There are 1580 Mi1s in FAFB (both sides), 1773 Mi1s in CNS (both sides), and 1654 Mi1s in BANC (extrapolating from 827 Mi1s in one proodread side). This is roughly a 12% span of Mi1 cell counts.

Altogether, although size differences cannot be ignored, they are not enough to account for the difference in synapse count.

## 3 Analysis

There are many differences in both the flies used and the reconstruction techniques that can confound any comparison. First, even flies of identical genetics, upbring, and sample preparation will be different. A good example is the Hemibrain, male CNS, and MANC flies. All were raised and prepared identically by the same researcher following the same protocals. Yet they have different synapse counts by up to 30%.

We examine a number of potential causes to see if they can explain the difference in synapse counts.

Is the difference the result of sexual dimorphism? This seems unlikely, as both nerve cords and brains of both sexes have been imaged with both SS-TEM and FIBSEM, and in every case the FIBSEM shows more synapses. In addition, we consider the cell Mi1, which shows more synapses in FIB-SEM than in FAFB. In this case, however, there is a third sample, FIB19 (Shinomiya et al. 2019), which is female like FAFB but imaged in FIB-SEM, like the Optic Lobe of the CNS. This is an excellent test case for checking for sexual dimorphism, as the flies are of different sexes, but both samples were raised similarly, imaged with similar FIB-SEM technology at similar resolution, and used similar synapse detection software. When we compare the outputs of the 1773 Mi1 cells in both optic lobes of the male CNS to the seven central columns of Mi1 (the columns most completely reconstructed) in the female FIB19, we find the counts are identical within the limits of the small sample size of FIB19. We therefore believe that Mi1 is not sexually dimorphic, and the difference in synapse counts between FAFB and CNS, as shown in Fig. 4, cannot be attributed to sexual dimorphism.

**Figure 4:**
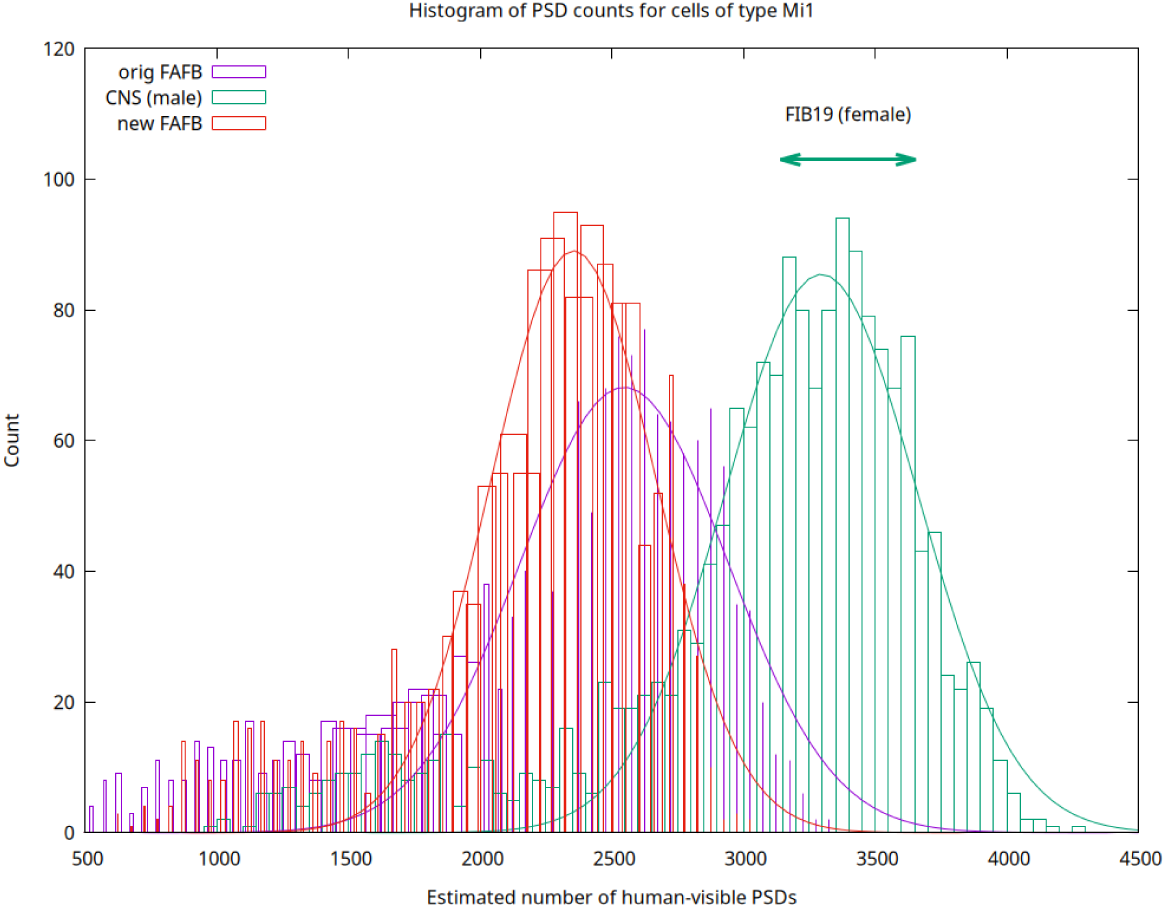
Comparison of synapse counts for Mi1 in three samples. For FAFB, FAFB2, and CNS the full distributions are shown over the thousands of cells in each sample. In contrast, FIB19 has only 7 fully reconstructed Mi1s, and hence is shown as a range (3398 ± 256) rather than a histogram. FIB19 is much more consistent with CNS (same imaging, different sexes) than it is with FAFB (same sex, different imaging).

Could the additional synapses found with FIB-SEM be an artifact (“hallucinated”) by the machine learning algorithms used to detect them? This seems unlikely, as they are well correlated with synapses already known to be present, as shown in Figures 1 and 3. Furthermore, this can be tested using the cell’s own opinion of whether the extra PSDs are real. In the hemi-brain sample, imaged by FIB-SEM (Scheffer et al. 2020) mitochondria have been identified in addition to synapses. As Fig 5 from (Rivlin et al. 2024) shows, as the putative PSD count increases, the nearest mitochondrion is both closer and bigger. The straightforward explanation is that these additional PSDs are real, and consuming energy.

**Figure 5:**
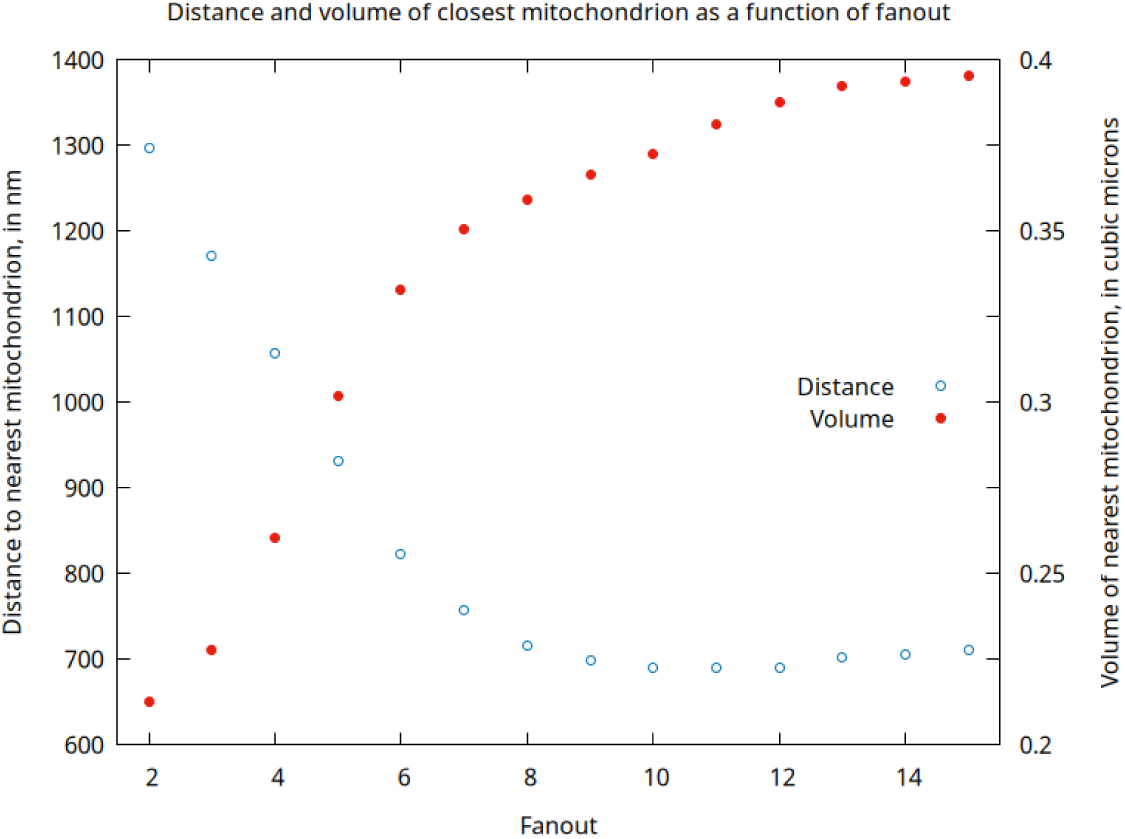
Plot of nearest mitochondrion distance and size as a function of synapse fanout, for hemibrain data set.

Could the differences be due to differing sample preparation? Methods for sample preparation have improved between the first and the most recent reconstructions(Lu et al. 2022, Pang & Xu 2023). (The FAFB sample, though reconstructed in 2024, was prepared and imaged many years earlier(Zheng et al. 2018).) In addition, sample preparation neccessarily differs between SS-TEM and FIBSEM due to the unique requirements of each technique. For example, SS-TEM requires embedding in a plastic that slices well, whereas FIBSEM needs a plastic that mills evenly. So for the largest examples, it is difficult to separate the effects of sample preparation and imaging.

However, there is evidence that sample preparation is not the cause of the observed differences. The single and seven column reconstructions (1C and 7C) differed in their imaging, 1C using SS-TEM and 7C using FIB-SEM. But the samples were prepared identically, specifically for the purpose of comparing the two results(Takemura et al. 2015). The FIBSEM reconstruction showed roughly 1.5x more synapses, consistent with the results of newer and larger reconstructions. We conclude that while sample preparation differences likely contribute to differences in synapse counts, they are a small contributor compared to other effects.

This leaves imaging techology as a suspect. In all cases but one where a sizeable circuit was reconstructed with both SS-TEM and FIB-SEM, a significantly higher number of synapses were found in the FIB-SEM sample (See Table 4). The ratios ranging from 0.89 (from one since superceded synapse detection) to 1.95, with a mean of 1.44 (geometric mean of 1.49). Roughly the same difference is noted in smaller reconstructions of individual areas. Takemura (Takemura et al. 2015) reported roughly 1.5x more synapses found per neuron in the 7 column FIB-SEM when compared to the SS-TEM single column, even though both were prepared similarly.

The only major exceptions to this trend occur in Schlegel et al. (2024), Extended Data Figure 2(e), where three brain regions appear to have significantly fewer synapses in the hemibrain when compared to FAFB. However, closer inspection shows these three regions (CAN(R), EPA(L), and LO(R)) are all truncated in the hemibrain sample but not in FAFB, so the lower synapse count is to be expected. This difference does not appear in our analysis, where we restrict comparisons to compartments fully contained in both reconstructions.

It is also suggestive, but again not proof, that if lesser Z resolution is part of this effect, this is consistent with the BANC and FANC samples (45 nm slices) finding somewhat fewer synapses than the FAFB sample (40 nm slices).

Overall, the most parsimonious explanation of synapse count differences is the greater ability of FIB-SEM to see synapses, in particular those in the direction perpendicular to the cutting plane (see Appendix 2). This explains both why there are more total connections found in the FIB-TEM data for similar circuits and the anisotropic nature of the synapses found in SS-TEM, as compared to the isotropic nature in FIB-SEM.

Fortunately, it appears the missing synapses for each method are randomly selected from the set of all synapses. Therefore, sufficiently strong paths in one reconstruction will almost always appear in another. This has been demonstrated directly by comparing reconstructions of differing completeness from the same sample (Scheffer et al. (2020), figure 16), and by computing two different symapse sets from the same data, as in FAFB and FAFB2. It also appears to true for different samples with different sample preparation and imaging techniques. This means many analyses are unaffected by a less complete reconstruction. Strong paths remain strong paths, and while their absolute strength is unknown, for many analyses a constant scale factor has little effect.

Using completion rates and synapse counts, we can estimate the odds that a path of a given biological strength is detected at all, as well as the odds it is reported as a strong (≥ 5) connection. To do this we necessarily make a number of assumptions and approximations. Hence the numerical results should be treated with suspicion, but comparative results are more likely true.

- First, we assume the underlying flies are the same, and the differences in observed synapse count are due to sample preparation, slicing, imaging, and synapse detection software.
- We assume that even the examples that report the greatest number of synapses are not finding them all. We arbitrarily assign this a factor of 0.9.
- We assume the odds that a synapse is visible is shown by the comparisons of regions in common, as shown in Table 4.
- Assuming the visibility of each synapse is independent, we compute a binomial distribution of expected synapse counts.
- For each synapse count, we estimate the number of independent segments comprising that connection by assuming a constant number *N* of synapses between the same neurons per fragment. In reality this is a distribution strongly dependent on the details of the segmentation. We compensate for this to some extent by running our analysis with fragment sizes of 1-5. In general this makes little difference for reasonably strong connections of weight 20-50, so we show curves for N=2. One exception is in the case of low completeness (particularly BANC). Here the odds of seeing a strong connection as strong rise considerably when N=5, as mere weak connections are no longer possible. Therefore for BANC-C, we show the curves for both N=2 and N=5.
- We then use the connection percentage as shown in Table 6 to estimate the odds that each segment is connected.
- We then find the odds that (zero, one, two, more) segments are detected. This allows us to compute the odds that a connection is reported at all, and that it is reported as a strong connection.

**Table 6:** Completion rates. Field ‘How computed’ refers to the condition that must be met for a synapse to count towards the completion rate. BANC is included twice, both with and without the optic lobes, as these are known to be at most partially reconstructed.

| Data set | Comp. | How computed |
| --- | --- | --- |
| FAFB | 41.90% | Both sides in ‘neuron_annotations.tsv’ |
| FAFB2 | 40.95% | Remove autapses, then same as above |
| HB | 35.25% | Both sides marked as ‘Neuron’ |
| CNS | 40.10% | Both sides marked as ‘Neuron’ |
| MANC | 44.43% | Both sides marked as ‘Neuron’ |
| BANC | 13.67% | Both sides in ‘neurons.csv’ |
| BANC-C | 16.02% | Central BANC, no optic lobes ME, LO, or LOP |

Using these assumptions, the odds of missing a connection of strength *S* is shown in Fig. 6 and Tables 7 - 8. Two curves and tables are shown - one for the odds of seeing a connection at all, and one for considering a connection only if the reported strength is ≥ 5, as many analyses consider only such paths are reliably reported.

**Figure 6:**
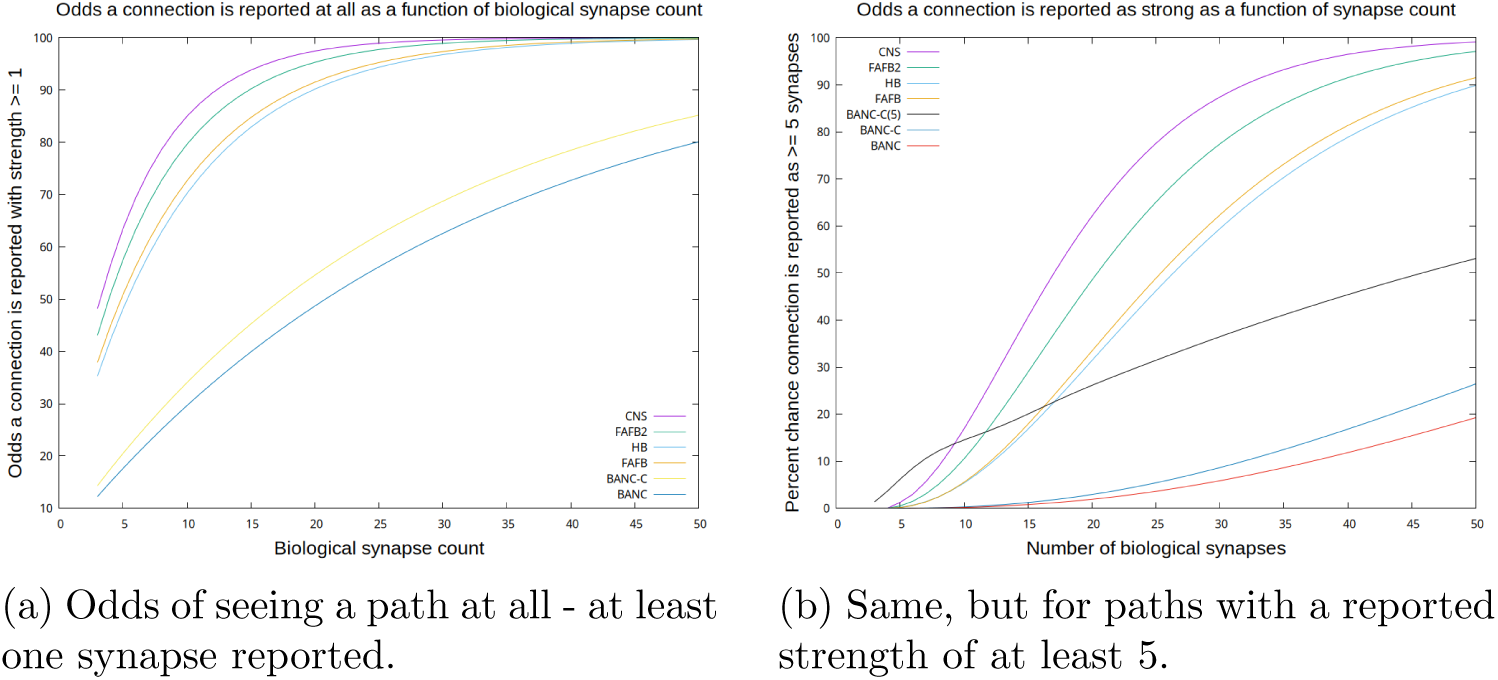
Odds of seeing a path of a given biological synapse count. Assumptions behind these estimates explained in text. Three BANC curves are shown: *N* = 2 for full BANC and for central brain, and *N* = 5 for the central brain only.

**Table 7:** Odds of seeing a path of specified strength at all.

| Strength | BANC | BANC-C | FAFB | FAFB2 | HB | CNS | MANC |
| --- | --- | --- | --- | --- | --- | --- | --- |
| 5 | 17.62% | 20.50% | 50.90% | 48.17% | 57.56% | 63.53% | 68.30% |
| 10 | 29.65% | 34.04% | 72.65% | 70.23% | 79.63% | 84.99% | 88.47% |
| 15 | 39.92% | 45.27% | 84.76% | 82.90% | 90.22% | 93.82% | 95.81% |
| 20 | 48.69% | 54.59% | 91.51% | 90.18% | 95.31% | 97.46% | 98.48% |
| 25 | 56.18% | 62.33% | 95.27% | 94.36% | 97.75% | 98.95% | 99.45% |
| 30 | 62.58% | 68.74% | 97.36% | 96.76% | 98.92% | 99.57% | 99.80% |
| 35 | 68.04% | 74.07% | 98.53% | 98.14% | 99.48% | 99.82% | 99.93% |
| 40 | 72.71% | 78.48% | 99.18% | 98.93% | 99.75% | 99.93% | 99.97% |
| 45 | 76.69% | 82.15% | 99.54% | 99.39% | 99.88% | 99.97% | 99.99% |
| 50 | 80.10% | 85.19% | 99.75% | 99.65% | 99.94% | 99.99% | 100.00% |

**Table 8:** Odds of a path of specified biological strength reported as strong, ≥ 5 synapses.

| Strength | BANC | BANC-C | FAFB | FAFB2 | HB | CNS | MANC |
| --- | --- | --- | --- | --- | --- | --- | --- |
| 5 | 0.00% | 0.01% | 0.16% | 0.20% | 0.49% | 1.25% | 1.70% |
| 10 | 0.18% | 0.29% | 5.54% | 5.37% | 10.52% | 16.90% | 21.71% |
| 15 | 0.77% | 1.21% | 17.89% | 16.82% | 28.94% | 40.69% | 48.87% |
| 20 | 1.90% | 2.92% | 33.45% | 31.42% | 48.52% | 62.11% | 70.55% |
| 25 | 3.60% | 5.41% | 48.90% | 46.22% | 65.08% | 77.57% | 84.39% |
| 30 | 5.85% | 8.64% | 62.36% | 59.44% | 77.45% | 87.42% | 92.20% |
| 35 | 8.62% | 12.49% | 73.16% | 70.34% | 85.97% | 93.23% | 96.27% |
| 40 | 11.83% | 16.82% | 81.36% | 78.85% | 91.52% | 96.47% | 98.27% |
| 45 | 15.41% | 21.51% | 87.32% | 85.22% | 94.99% | 98.20% | 99.22% |
| 50 | 19.26% | 26.43% | 91.53% | 89.85% | 97.10% | 99.10% | 99.65% |

One conclusion from these charts is that if an analysis depends on identifying strong paths as strong, then completeness is a critical need in reconstruction. Reconstructions with low completion rates (much less than 40%), at least according to the model proposed here, are quite likely to miss paths with biological strengths in the range of 20-50 synapses. (In fairness to the BANC community, their lower completeness was not driven by the sense that lower rates are sufficient, but instead by funding cuts preventing further increases in completeness (Alexander Bates, personal communication).)

On the other hand, comparing connectomes by their measured completeness can even provide a perverse incentive towards incomplete synapse detection. One of the major difficulties in *Drosophila* reconstruction is connecting the tiny fragments containing PSDs to larger segments whose identity is known. If fewer synapses are found, less of this demanding work is required to reach the same measured completeness.

## 4 Conclusions and future research

A completely unambiguous comparison of synapse detection rates between the *Drosophila* samples analyzed here is not possible from the data given, given the very small *N* and differences in the ways the flies were selected, raised, stained, and imaged. However, the weight of the evidence points to samples imaged with FIB-SEM detecting more synapses, typically by about 40%.

There are experiments that could settle the questions raised here. If the limiting factor is sample preparation, more recent reconstructions with improved sample preparation should yield more equal estimates of synapse count. Another possibility is that multi-color expansion microscopy could reliably label all Tbars and post-synaptic partners and resolve them so they might be counted. Then identically raised flies could be analyzed by various EM techniques to find the absolute synapse detection efficiency.

To determine if the resolution differences account for the synapse count differences, one possibility might be to image a volume at 4×4×4 nm resolution with FIB-SIM. Then the data could be summed two ways - into 8×8×8 nm cubes, and 4×4×40 nm slices. Then both synapse detection algorithms could be tried and the results compared with each other and ground truth.

If the problem is anisotropic resolution, there are a number of improvements that could be pursued. Perhaps humans aware of this particular configuration could do better, and create better ground truth. And even if humans cannot find these synapses, it is possible that machine learning can spot subtle signs that humans tend to miss, as is true for neurotransmitter ID(Eckstein et al. 2024).

## 5 Data Availability

Most of the data used in this paper is publicly available, with the rest obtained from the authors of the respective papers. The FAFB data, version 783, was downloaded from https://github.com/flyconnectome/flywire_annotations/tree/main and https://zenodo.org/records/10676866.

The data for the hemibrain, the optic lobe of the CNS, the FIB19 data set, MANC, and the full male CNS, can be found at https://neuprint.janelia.org. The CREMI challenge data set can be found at https://cremi.org/data/. The data for BANC can be found at https://codex.flywire.ai/banc. The full synapse data set for FAFB2 was provided by Sven Dorkenwald. The full synapse data set for FANC was provided by Jasper Phelps.

## 6 Acknowledgements

Much of this work was done as part of The FlyEM Project Team at Janelia Research Campus in collaboration with the Drosophila Connectomics Group, University of Cambridge, and the associated labs. In particular the provided pre-publication access to the male CNS data. Funding was provided by HHMI/FlyEM and the Wellcome Trust.

## 7 Appendix 1 - Differences between flies

The samples compared here are from different flies and sometimes different sexes. The flies were in many cases raised differently. For example, the FAFB fly was raised in the Janelia EM facility, and the CNS fly in the fly core facility. This resulted in the following differences, all of which are known to affect connectivity.

- The FAFB fly was female, and the CNS fly was male.
- The FAFB flies were raised in 24 hour light (sometime bright and sometimes dim, depending on the motion sensing), where the CNS flies were raised on a 12 hour on, 12 hour off schedule. Circadian rhythm is known to have an effect on synapse counts(Pyza & Meinertzhagen 1993).
- The temperature was different in the two facilities. The EM facilities are normally kept at 21-22*^◦^* C, whereas the fly core incubation area is kept at 24*^◦^* C (Zhiyuan Lu, personal communication). Temperature differences are known to significantly affect synapse count, at least in the optic lobe(Kiral et al. 2021) and olfactory system(Züfle et al. 2025), where a factor of two difference has been observed between flies raised at 18 and 25*^◦^* C. However, this is unlikely to account for the observed FAFB ⇔ FIB-SEM synapse count differences, as the temperature difference predicts more synapses in FAFB than CNS, but fewer are observed.
- The available food supplies were likely different. The FAFB sample has a higher count of Kenyon cells(Dorkenwald et al. 2024), which is a known effect of protein deprivation during development(Lin et al. 2013).
- The FAFB fly was sampled 7 days post-eclosion, where the CNS and Hemibrain flies were sampled at 5 days. Age is known to affect synapse count(Jay et al. 2025).

The SS-TEM and FIB-SEM flies are embedded in different plastics, each optimized for the sectioning method used. The imaging resolutions used were different - SS-TEM uses roughly 4×4×40nm, and FIB-SEM gives about 8nm isotropic. The segmentation and annotation software was different. The synapse detection software was quite different. The fly has polyadic synapses, where a single pre-synaptic density (commonly called a TBar) is presynaptic to multiple post-synaptic densities (PSDs). We refer to the number of PSDs for a single TBar as the *fanout*. In the FIB-SEM reconstructions this structure is explicit, with a single TBar linked to multiple partners. In the FAFB reconstruction the structure is implicit. Each PSD is linked with its own point in the pre-synaptic neurite, but the pre-synaptic points are not combined, nor identified, as TBars.

## 8 Appendix 2: Potential causes

In this section we examine some differences in the synapses found in the original FAFB synapses, as opposed to those found in the HemiBrain.

The main difference in the imaging is the relatively low Z resolution of SS-TEM, especially in contrast to high resolution (4 nm) in X and Y. This leads to the question of whether synapses are detected equally well in all orientations. To test this, we calculate a vector from the pre to post points of each synapse. The directions are mapped onto a sphere surrounding the origin. If the synapses are isotropically oriented, and the synapse detection non-biased, we should see an even distribution around this sphere. When we view the sphere for FIB-SEM this is exactly what we see, as shown in Fig. 7, where we view the sphere from the perpendicular to, and along, the Z plane.

**Figure 7:**
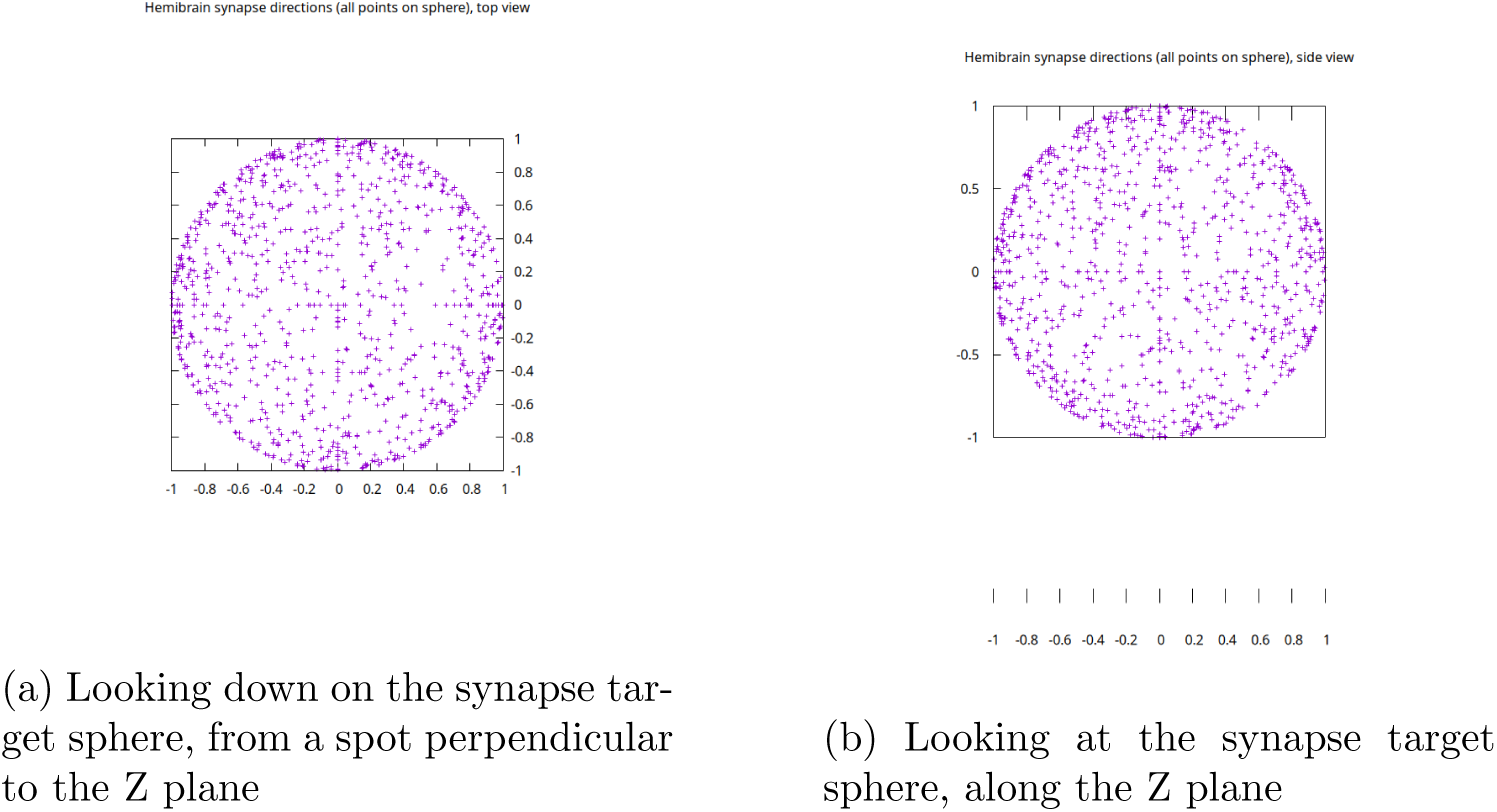
In FIB-SEM, the synapses and their detection are roughly isotropic.

In FAFB, however, the pattern looks very different. The synapses are concentrated around in the plane of the Z-cut, and many fewer are found in the direction perpendicular to the cut. See Fig. 8.

**Figure 8:**
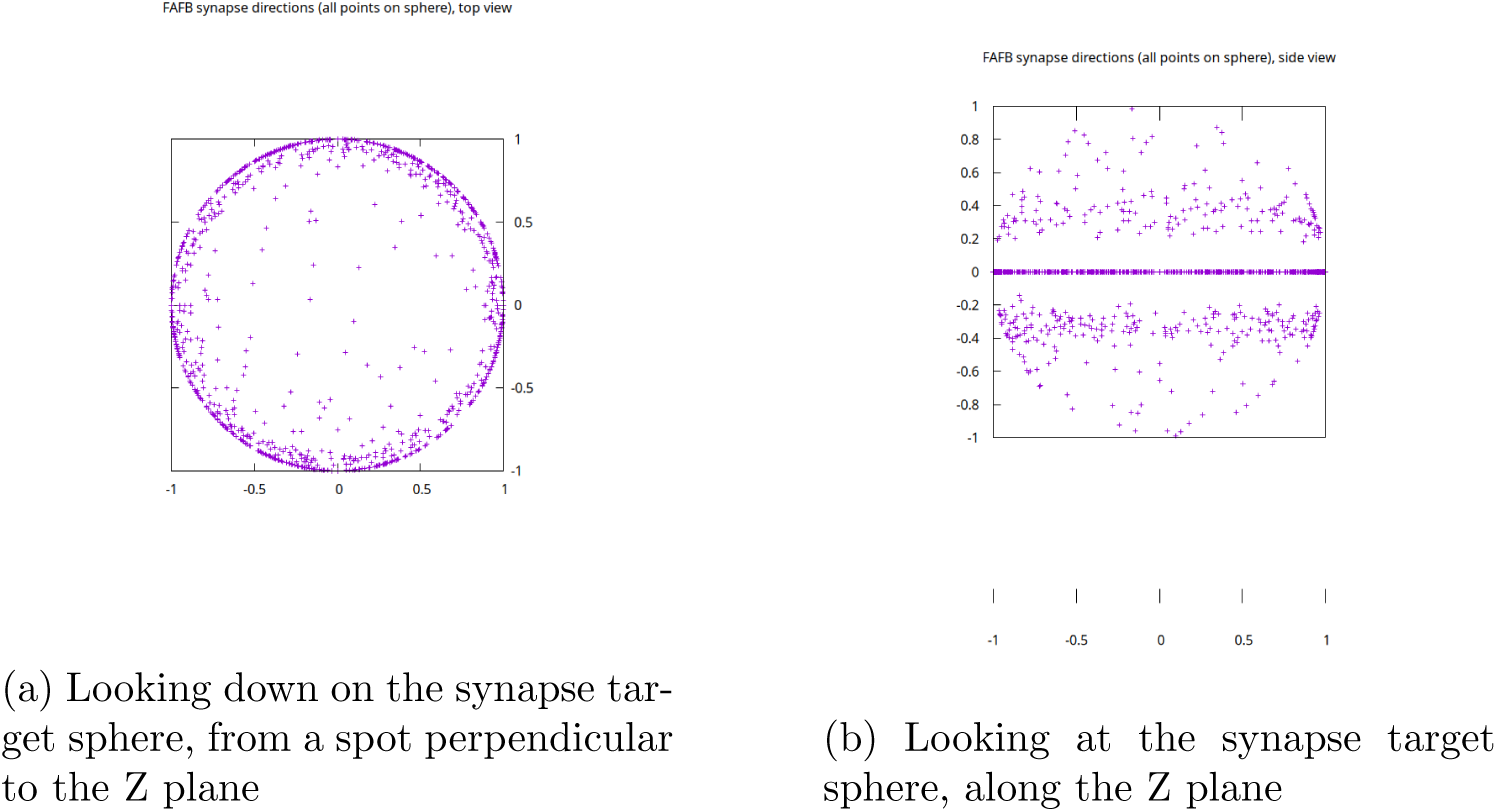
In SSTEM, the synapses and their detection are not isotropic. Many more synapses are detected along the Z-cut plane than are detected perpendicular to it.

We can quantify how anisotropic this pattern is. We can divide the sphere of synapse detection into three bands - within 19.5*^◦^* of the equator, between 19.5*^◦^* and 41.5*^◦^* from the equator, and above 41.5*^◦^*. These angles were picked so the areas on the sphere are equal, and hence the percentage of synapses should be as well if they are isotropic. The measurements for the Ellipsoid Body (EB) in FAFB and hemibrain are shown in Table 9. Other regions, and random sampling of the synapse sets, give similar results (not shown).

**Table 9:** Statistics of synapse count by angle from cut plane. Data is from the region EB in FAFB and the hemibrain (HB). Ground truth is the combined ground truth for the FAFB synapse prediction (EB, LH, and MB). *θ* is the angle of the pre→post vector from the cut plane. The FIB-SEM distribution is close to isotropic; the SS-TEM distribution and the ground truth are not.

| Data set<br>name | Band 1<br>$\theta < 19.5^\circ$ | Band 2<br>$19.5^\circ < \theta < 41.5^\circ$ | Band 3<br>$\theta > 41.5^\circ$ | Total |
| --- | --- | --- | --- | --- |
| HB count<br>HB % | 258135<br>35% | 255990<br>35% | 215173<br>30% | 729298 |
| FAFB count<br>FAFB % | 350934<br>78% | 86830<br>19% | 10128<br>2.3% | 447892 |
| Ground truth<br>percent | 62513<br>71.5% | 16325<br>18.7% | 8650<br>9.9% | 87488 |

Just because they synapses are *reported* anisotropically does not necessarily mean they were *detected* anisotropically. The distinguishing points for pre- and post-synapses are not unique, and could be picked differently by either a person or an algorithm. This does not affect the count of synapses but could affect the angular bin in which they fall. In particular it is possible that the FAFB synapses are more commonly reported as low-angle when compared to HB synapses. This could be due to the quantization of possible Z values, or because single plane annotations are easier to annotate for humans, an attribute that could be learned from training data (see below).

This could explain why when considering the absolute number of synapses detected, we see that FAFB reports more low-angle (with respect to the cutting plane) synapses then the hemibrain, but fewer high-angle synapses. The two numbers become equal near 30*^◦^*. A plausible (but certainly not the only) interpretation is that both algorithms have similar ability to detect near in-plane synapses, but FAFB tends to report them at lower angles.

As mentioned above, an obvious question is what pattern is found when a human detects synapses in SS-TEM data. For this we can look at the training data for the synapse detector used in FAFB(Buhmann et al. 2021). This data is publicly available as part of the CREMI challenge. Like the automatically detected synapses, the human detected synapses are far from isotropic, as shown in Fig. 9. This anisotropy has the same signature as the FAFB data - it is strongly biased towards in-plane synapses. The same holds true of the full training set used in the detection of FAFB synapses - see Table 9.

## 9 Appendix 3: Region mapping for nerve cord of BANC and FANC

**Figure 9:**
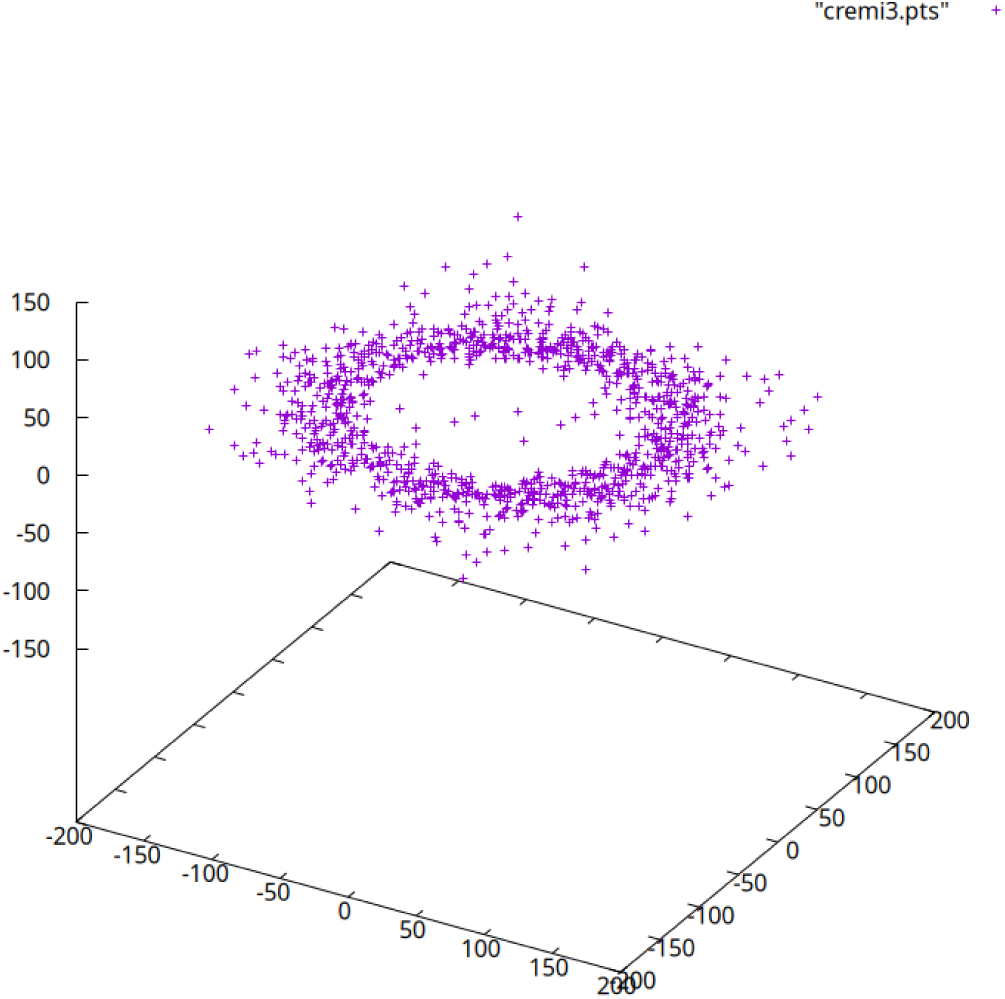
Orientation of manually detected synapses in the CREMI training data. Data is from CREMI Dataset C. Sets A and B are similar but not shown.

**Table 10:** Mapping from FANC to CNS/MAC region names.

| FANC | CNS/MANC | BANC | CNS/MANC |
| --- | --- | --- | --- |
| AMNp_L | Ov(L) | AMNP | Ov |
| AMNp_R | Ov(R) | ABDNM | ANm |
| HTct_L | HTct(UTct-T3)(L) | HTct | HTct(UTct-T3) |
| HTct_R | HTct(UTct-T3) |  |  |
| NTct_L | NTct(UTct-T1)(L) | NTct | NTct(UTct-T1) |
| NTct_R | NTct(UTct-T1)(R) |  |  |
| WTct_L | WTct(UTct-T2)(L) | WTct | WTct(UTct-T2) |
| WTct_R | WTct(UTct-T2)(R) |  |  |
| ProNm_L | LegNp(T1)(L) | ProNM-T1 | LegNp(T1) |
| ProNm_R | LegNp(T1)(R) |  |  |
| MesoNm_L | LegNp(T2)(L) | MesoNM-T2 | LegNp(T2) |
| MesoNm_R | LegNp(T2)(R) |  |  |
| MetaNm_L | LegNp(T3)(L) | MetaNM-T3 | LegNp(T3) |
| MetaNm_R | LegNp(T3)(R) |  |  |
| mVAC_T1_L | mVAC(T1)(L) | mVAC | mVAC(T1) |
| mVAC_T1_R | mVAC(T1)(R) |  |  |
| mVAC_T2_L | mVAC(T2)(L) |  |  |
| mVAC_T2_R | mVAC(T2)(R) |  |  |
| mVAC_T3_L | mVAC(T3)(L) |  |  |
| mVAC_T3_R | mVAC(T3)(R) |  |  |

